# Anti-Diabetic Effects of *Nigella sativa and Cichorium intybus* in Animal Experimental Model

**DOI:** 10.1101/2025.02.03.636360

**Authors:** Muhammad Kashif, Amar Nasir, Muhammad Khizer Aziz, Aziz ur Rehman, Tom Vincent, Jean-Paul Gonzalez, Usman Waheed

## Abstract

Diabetes mellitus is a chronic metabolic disease in humans associated with severe complications with a significant impact on the health, life expectancy, and lifestyle of patients.

The present study has been designed to examine the anti-hyperglycemic efficacy of two medicinal plants - *Nigella sativa* seeds and *Cichorium intybus* leaves - and their combined effects on body weight and hematological/serological parameters in alloxan-induced diabetic rabbits.

For this purpose, 30 adult rabbits were divided into six equal groups of five rabbits each. Group I was kept as the negative control, while all other groups received a single intravenous injection of alloxan monohydrate to induce diabetes: Group II as positive control; Group III received metformin hydrochloride; Group IV was fed powdered *C. intybus* plant leaves; Group V was given *N. sativa* aqueous seeds extract; and Group VI was fed a combination of *C. intybus* and *N. sativa*. Fasting blood glucose levels and body weight of rabbits were observed at 0 days (before treatment) to Day 28 and regularly monitored during the four-week experimental trial. Biochemical analysis, serum insulin level, and hematological parameters were measured at the end of the experiment.

Results showed that *C. intybus, N. sativa*, and their mixture significantly (*p* < 0.05) decreased the blood glucose levels, urea and creatinine levels, and enzyme levels in the blood. At the same time, body weight gain, red blood cell (RBC) count, white blood cell (WBC) count, and packed cell volume (PCV) % were increased in diabetic rabbits as compared with positive control rabbits. It was concluded that these plants have potential hypoglycemic effects and reduced diabetic complications in alloxan-induced diabetic rabbits. Further studies are needed to understand their potential for antidiabetic remedy.

## BACKGROUND

Diabetes mellitus is a carbohydrate metabolic disorder that is characterized by the body’s response to generate insulin and maintain suitable blood glucose levels. It is a major health concern associated with many functional and structural metabolic complications. Diabetic complications include ketogenesis, gluconeogenesis, and increased risk of heart attacks and strokes (Samoo et al., 2018). According to the International Diabetes Federation, approximately 194 million people had diabetes in 2003, which will rise to 333 million in 2025. There are two major types of diabetes: Type 1 (TD1, insulin-dependent, or juvenile diabetes mellitus) and Type 2 (TD2, non-insulin-dependent diabetes mellitus). It is estimated that more than 90% of all diabetic patients have T2D (Islam and Choi 2008). Diabetes mellitus is present in both developed and underdeveloped countries in the world. In low- and middle-income countries, the number of diabetic patients will increase from 84 million to 228 million, whereas these numbers will rise from 21 million to 72 million patients in high-income countries (HICs). It is assumed that at the end of 2025, about 70% of the diabetic ratio belongs to HICs (Ashraf et al., 2011).

Diabetes mellitus is the principal cause of morbidity and mortality in the world (El-Kott et al., 2010). According to the International Diabetes Federation (IDF), diabetes affects 2.8% of total world population. It is also a serious problem in the Indo-Pak region. Pakistan National Diabetes Survey (2016-2017), showed approximately 35.5 - 37.5 million people are affected by diabetes in Pakistan. Diabetes mellitus and its complications are primarily caused by hyperglycemia and lifestyle-related issues (Venkatesh et al., 2003; Mukherjee et al., 2006). Herbal therapies for diabetes treatment have been used for decades in TD1 and TD2 patients.

In Pakistan, it is projected that the number of patients suffering from diabetes will rise from 4.3 million, in 1995 to 14.5 million, in 2025. The pattern of provincial community diabetes distribution in Pakistan shows Sindh accounting for 33.5%, followed by Punjab with 31.3%, Baluchistan with 28.2%, and Khyber Pakhtunkhwa with 13.2%. These numbers apply to about 27.4 million people aged twenty and over, based on a total population of 207.8 million. If the current situation persists, the highest incidence of diabetes worldwide is expected to be in Pakistan. For the health concern community and public health care policymakers, this current diabetes situation presents a tremendous challenge (Abo et al., 2008).

In recent years, the use of natural substances has been increased among the general public for multiple diseases not only because of their easy availability to health care professionals, cost-effectiveness, and varieties, but also because of the awareness that natural products have fewer side effects than synthetic medicines. The use of herbal plants mostly does not have drugs like mode of action or adverse hazards (Karmi et al., 2015). Various natural products have been tested for their possible antidiabetic effects (Bamosa, 2015).

Black cumin seed (*Nigella sativa*) and Chicory or Kasni (*Cichorium intybus)* are herbs commonly used to treat different diseases in the world. *N. sativa* is a seasonal herb, that belongs to the *Ranunculaceae* family, often grown in the Indian Subcontinent, Africa, and the Middle East (Benhaddou-Andaloussi et al., 2008). *N. sativa* seeds are sometimes practiced as spices but are often commonly used in traditional medicines in many of these countries (Farkhondeh et al., 2017). This plant has historically been mentioned in Islamic culture because of having dramatic healing properties (Al-Bukhari, 1976). Conventional food ingredients, such as garlic, ginger, and bitter gourd have been used extensively that can decrease blood glucose, hepatic gluconeogenesis, intestinal glucose intake, body weight and animate the insulin levels in diabetic patients (Imo, 2019). It has been demonstrated that diabetic-induced rats and hamsters treated with Black seed *(N. sativa)* seeds and their components did not show any adverse effects (Mathur et al., 2011). Black seed has also been used against different human diseases such as hepatitis, diarrhea, fever, cough, and tapeworm disease (Hamdan et al., 2019). It is reportedly beneficial for immune response (El-Rabey et al., 2017). Ultimately, *N. sativa* has many beneficial effects, including antidiabetic effects, cardiovascular, anti-cancer, and renal effects, as well as anti-inflammatory, anti-parasitic, anti-asthmatic, and anti-hypertensive effects. In addition, it is also used to cure various disorders like respiratory issues, diarrhea, and skin disorders (Børve and Børve, 2020). *N. sativa* oil extract has greatly improved clinical symptoms in patients with allergic disorders such as bronchial asthma, allergic rhinitis, and atopic eczema (Das et al., 2016).

*C. intybus* is widely cultivated in a variety of temperate regions around the world, including South Africa and the Indian Subcontinent. It is used for the treatment of AIDS, cancer, diabetes, splenitis, tachycardia, gallstones, gastroenteritis, sinus issues, cuts, and bruises as well as the energizing effect on the digestive tract and liver (Ghamarian et al., 2012; Abdel-Rahim et al., 2016). This plant contains several active ingredients including caffeic acid derivatives, flavonoids, polyphenols, and prebiotic inulin that have a potential for many properties such as anti-inflammatory, sedative, anticancer, cardiovascular, gastroprotective, hepatoprotective, reproductive, antioxidant, analgesic, hypolipidemic, anthelmintic, and antimicrobial effects (Abdel-Daim and Ghazy 2015; Rafiqi et al., 2019, Sakihama et al., 2002). The water and alcoholic extract of the plant revealed hepatoprotective effects in experimentally-induced liver damage in rats and an antibacterial activity against *Staphylococcus aureus* in the aqueous seed extract (Ahmed 2009; Mitra et al., 1998).

Our study has been designed to examine the effects of *N. sativa* and *C. intybus* on blood glucose levels, serum insulin levels, body weight, hematological parameters, liver and kidney function enzymes, in alloxan-induced diabetic rabbits.

## MATERIALS AND METHODS

### Chemicals

Alloxan monohydrate (MP Biomedicals, LLC, Parc innovation, BP 50067 IIIkirch, France) was used for induction of diabetes in rabbits; Tab Metformin 5 gm from Sigma Chemicals Co. (St. Louis, Mo, USA) was used as standard drug for comparison of antidiabetic potential of experimental plants; Glucometer from Capricorn Scientific (Sujeong-gu, Seongna, South Korea) used for measuring blood glucose level); spirit and alcohol were used as antiseptic and were purchased from the local market in Pakistan.

### Collection and extraction of plant material

*N. sativa* and *C. intybus* plants were purchased from the National Agricultural Research Center (NARC) in Islamabad, Pakistan. *N. sativa* seeds were washed, dried, and crushed into powder using a mortar & pestle. Aqueous extract of *N. sativa* seeds (50 g in distilled water 1000 ml for 2-4 hours) was obtained by using a Soxhlet apparatus. The solvent was removed from the extract, using a hot air oven (65°c for 24hrs). *N. sativa* seed extract was collected and used for the experiment.

*C. intybus* leaves dried for 21 days, then finely powdered and stored in airtight glass containers; an extraction procedure similar that of the *N. sativa* seeds was then used (Hassan & Sudomova, 2020; Kahkonen et al., 1999).

### Phytochemical analysis of plants

Phytochemical analyses of *C. intybus* and *N. sativa* plants were performed in the WTO laboratory (World Trade Organization Accredited) at the University of Veterinary and Animal Sciences, Lahore. The quantity of alkaloids was observed through a spectrophotometer at 565 nm (Mitruka & Rawnsley, 1977), the quantity of phenol was determined by a reagent protocol method known as Folin-Ciocalteu (Abdullah et al., 2017, quantity of flavonoids was measured by a spectrophotometer at 415 nm (Zafar & Ali, 1998) and the number of tannins was evaluated by the Folin Denis reagent and the concentration of saponins was measured by the gravimetric method (Akhtar et al., 2020).

### Experimental Animals

Thirty adult male Dutch rabbits (mean weight = 1.25 kg) were purchased from the local market of Jhang City (Punjab Pakistan) and kept in the experimental animal house of the Department of Clinical Sciences, College of Veterinary & Animal Sciences, sub-campus Jhang. The rabbits’ maintenance followed the “Principles of Laboratory Animal Care’’, and were fed on a normal diet and cleaned, filtered drinking water throughout the experiment. The study plan was dually approved by the Ethics Committee of the University of Veterinary & Animal Sciences, Lahore (No. CVAS/13545 dated 07-01-2021) and by the Directorate of Advanced Studies of the same university (DAS/536 dated 15-06-2021).

### Diabetes Induction

Rabbits were kept fasted for 14 hours before induction of diabetes by a single intravenous injection of alloxan monohydrate (MP Biomedicals, LLC, Parc innovation, BP 50067 IIIkirch, France) with 80 mg/kg body weight after dissolving in normal saline. After induction of diabetes, animals were given 10 ml of 10% glucose solution to avoid death from hypoglycemic shock. After three days, the overnight fasting blood glucose levels were calculated using a glucometer (Accu Chek Performa^®^). The rabbits with fasting blood sugar levels ≥ 300 mg/dl, were considered diabetic and used for the experiment (Ahmed et al., 2014).

### Experimental design and treatment protocol

The selected rabbits were divided equally into six groups of five, and were treated according to the study protocol for 28 days. The study protocol for each group was as follows: Group I was kept as Negative control (with no diabetes induction and no treatment was given); Group II was kept as Positive control (diabetes induction and no treatment); Group III (with diabetes induction) was treated orally with 500 mg/kg Metformin; Group IV (with diabetes induction) was orally fed with 500 mg/kg *C. intybus* leaves powder; Group V (with diabetes induction) was orally fed with 300 mg/kg of *N. sativa* aqueous seeds extract; and, Group VI (with diabetes induction) was orally fed with a combination of 250 mg/kg *C. intybus* and 150 mg/kg *N. sativa* in one compound.

### Determination of fasting blood glucose levels and weight gain

Fasting blood glucose levels were calculated using a glucometer (Accu Chek Performa ®). Glucose levels were observed at from day “0” (before treatment), then every day from day 6 to day 28 days regularly after one week in the experiment. The body weight of all animals was calculated on a digital weighing balance at day 0 and a weekly interval in the study protocol.

### Blood sampling for Hematological examination

On day 28, the final day of the experiment, rabbits were fasted for an overnight period. Blood samples were obtained from the marginal ear vein using a sterile syringe and needle. These samples were transferred to vacutainers (BD Vacutainer® spray-coated K2 EDTA), for further analysis of insulin concentration and biochemical values. A micro-hematocrit centrifuge was used to determine the packed cell volume (PCV). These samples were diluted with diluting fluid in WBC and RBC counting pipettes, and a hemocytometer was used to count the individual cells. Giemsa staining was also used for RBC count (Rafiq et al., 2009).

### Determination of Insulin level

Three ml of blood samples were utilized for determining serum insulin serum concentration, and was done by radioimmunoassay using insulin enzyme-linked immunosorbent assay and immunoreactive insulin (free insulin + insulin bound to anti-insulin antibodies). The values of experimental groups were compared with the control groups (Meral et al., 2004).

### Biochemical Analysis

Blood samples of 2.5 ml were collected using non-heparinized tubes for harvesting serum and stored until used for biochemical analysis. Fasting serum enzyme activities, alanine transaminase (ALT), alkaline phosphatase (ALP), aspartate transaminase (AST), creatinine, and urea levels were measured by their respective kits using an automatic chemistry analyzer according to the manufacturer’s instructions.

### Statistical Analysis

The data was represented as means ± one standard deviation (SD). The statistical analysis was conducted with one-way ANOVA analysis and Tukey’s post hoc test (BRM-SPSS software, version 24).

## RESULTS

### Phytochemical Analysis of *Cichorium intybus* and *Nigella sativa* Plants

The key contents identified in the *N. sativa* plants included flavonoids (11.2±0.25 mg/g), saponins 68.55±0.40mg/g), alkaloids (15.21±0.30 mg/g), and tannins (13.60±0.20 mg/g) whereas *C. intybus* contains flavonoids (120.5±0.55 mg/g) alkaloids (17.40±0.10 mg/g), saponins (170.20±0.60 mg/g) and tannins (25.40±0.70 mg/g) (Table 1).

**Table 1:**
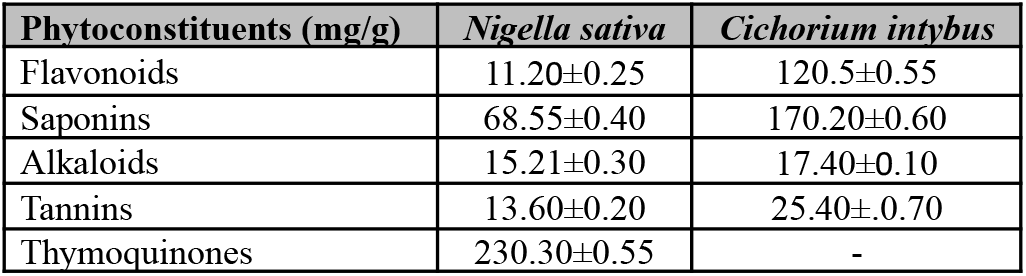
Quantitative Phytochemical Analysis of Aqueous Extracts of Studied Plants.

| Phytoconstituents (mg/g) | *Nigella sativa* | *Cichorium intybus* |
| --- | --- | --- |
| Flavonoids | 11.20±0.25 | 120.5±0.55 |
| Saponins | 68.55±0.40 | 170.20±0.60 |
| Alkaloids | 15.21±0.30 | 17.40±0.10 |
| Tannins | 13.60±0.20 | 25.40±0.70 |
| Thymoquinones | 230.30±0.55 | - |

### Effects of *Nigella sativa* and *Cichorium intybus* aqueous extract on fasting blood glucose levels in alloxan-induced diabetic rabbits

Mean blood glucose values were decreased gradually from (308±5.01) at 0 days (before treatment) and (268.6±4.45) on the 28^th^ day in *N. sativa*-treated diabetic animals. while these values were (308.4±4.96) at 0 days (before treatment) and (270.6±4.12) on the 28^th^ day in *C. intybus*-treated diabetic animals. The result shows that the mean values of blood glucose levels were significantly (*p* < 0.05) decreased in different treatment groups as compared with the positive control group. However, a slight difference in these values was shown among different treatment groups as compared with standard drugs. Moreover, the hypoglycemic effects of plant material were almost the same (Table 2).

**Table 2.**
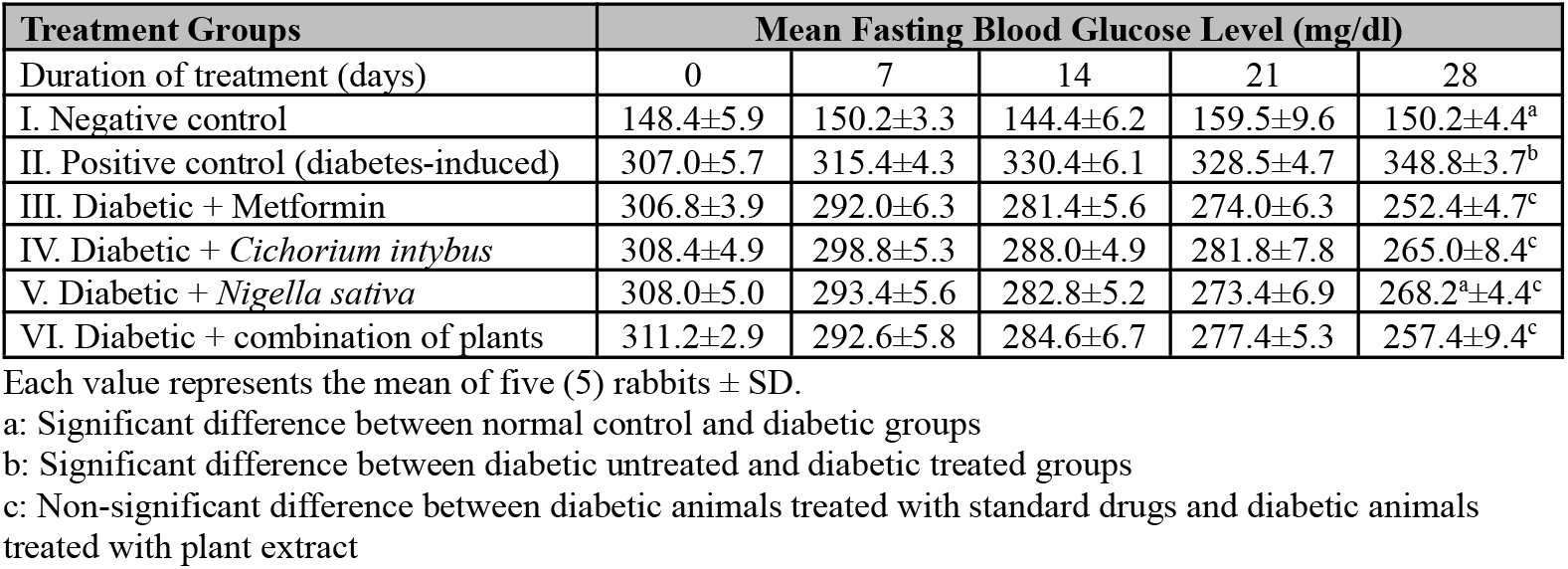
Effects of Nigella sativa and Cichorium intybus aqueous extract on fasting blood glucose levels in alloxan-induced diabetic rabbits.

| Treatment Groups | Mean Fasting Blood Glucose Level (mg/dl) |  |  |  |  |
| --- | --- | --- | --- | --- | --- |
| Duration of treatment (days) | 0 | 7 | 14 | 21 | 28 |
| I. Negative control | 148.4±5.9 | 150.2±3.3 | 144.4±6.2 | 159.5±9.6 | 150.2±4.4<sup>a</sup> |
| II. Positive control (diabetes-induced) | 307.0±5.7 | 315.4±4.3 | 330.4±6.1 | 328.5±4.7 | 348.8±3.7<sup>b</sup> |
| III. Diabetic + Metformin | 306.8±3.9 | 292.0±6.3 | 281.4±5.6 | 274.0±6.3 | 252.4±4.7<sup>c</sup> |
| IV. Diabetic + *Cichorium intybus* | 308.4±4.9 | 298.8±5.3 | 288.0±4.9 | 281.8±7.8 | 265.0±8.4<sup>c</sup> |
| V. Diabetic + *Nigella sativa* | 308.0±5.0 | 293.4±5.6 | 282.8±5.2 | 273.4±6.9 | 268.2<sup>a</sup>±4.4<sup>c</sup> |
| VI. Diabetic + combination of plants | 311.2±2.9 | 292.6±5.8 | 284.6±6.7 | 277.4±5.3 | 257.4±9.4<sup>c</sup> |

Each value represents the mean of five (5) rabbits ± SD.
a: Significant difference between normal control and diabetic groups
b: Significant difference between diabetic untreated and diabetic treated groups
c: Non-significant difference between diabetic animals treated with standard drugs and diabetic animals treated with plant extract

### Effects of *Nigella sativa* and *Cichorium intybus* aqueous extract on serum insulin levels in alloxan-induced diabetic rabbits

Insulin level was measured on Day 28 at the end of the experiment. The current study result showed that *C. intybus* and *N. sativa*, or their combination treatment significantly (*p* < 0.05) increased the serum insulin level in diabetic groups as compared with the positive control group. Meanwhile, the effect of plant material to increase serum insulin levels in diabetic groups was comparable to metformin treated diabetic group (Table 3).

**Table 3.** Effects of *Nigella sativa* and *Cichorium intybus* Aqueous Extract on Serum Insulin Levels in Alloxan-Induced Diabetic Rabbits.

| Treatment Groups | Mean Fasting Blood Glucose Level (μ/ml) after 4 Weeks |
| --- | --- |
| I. Negative control | 3.35±0.97 |
| II. Positive control (diabetes-induced) | 8.12±1.43 |
| III. Diabetic + Metformin | 7.18±1.07 |
| IV. Diabetic + *Cichorium intybus* | 7.98±1.34 |
| V. Diabetic + *Nigella sativa* | 6.79±1.56 |
| VI. Diabetic + combination of plants | 8.01±1.63 |

Each value represents the mean of 5 rabbits ± SD.

### Effects of *N. sativa* and *C. intybus* aqueous extract on body weight gain in alloxan-induced diabetic rabbits

The rabbit body weight was measured before and after diabetes induction. The mean values of body weight gain were increased gradually from (1.17 ± .062) at Day 0 (before treatment) and (1.42 ± .027) at Day 28 day in *N. sativa-*treated diabetic animals. while these values were (1.14 ± .094) at 0 days (before treatment) and (1.37 ± .036) on the 28^th^ day in *C. intybus-*treated diabetic animals. The current study result shows that the mean body weight gain values of treatment groups were significantly (*p* < 0.05) increased as compared with the positive control group. A slight difference in these values was shown in *N. sativa, C. intybus*, or their mixture-treated diabetic animals as compared with standard drugs (Table 4).

**Table 4.**
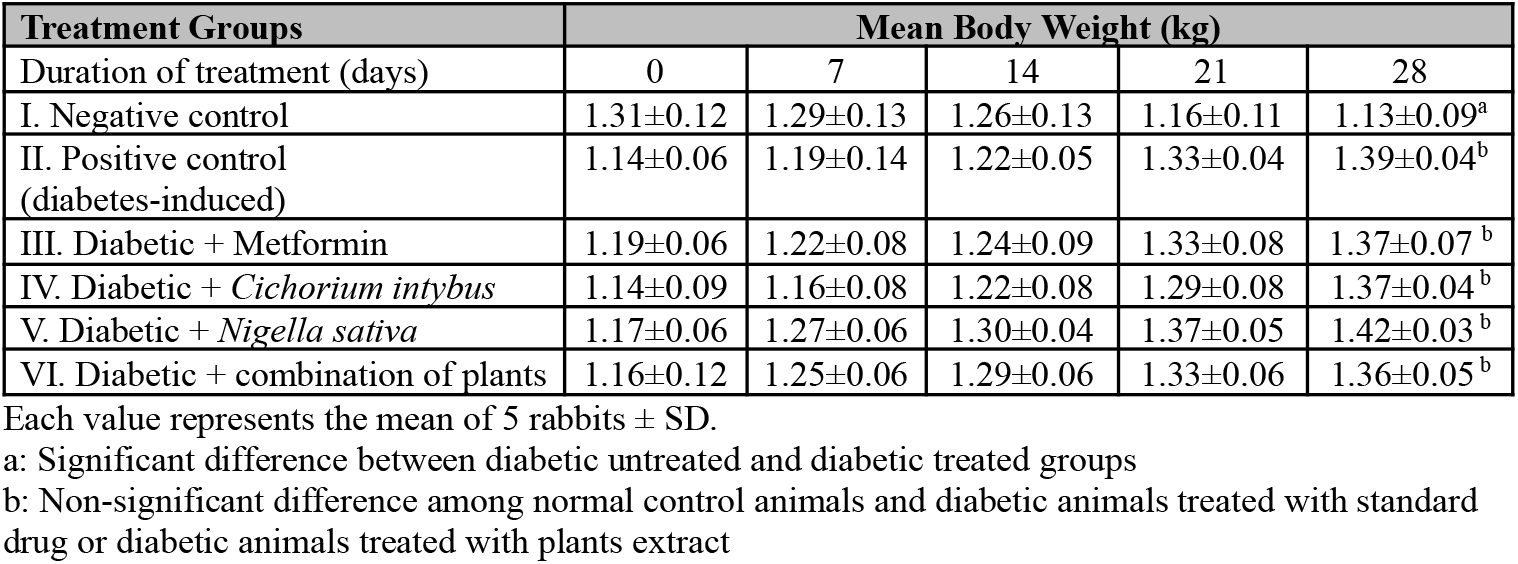
Effects of *Nigella sativa* and *Cichorium intybus* Aqueous Extract on Body Weight Gain in Alloxan-Induced Diabetic Rabbits.

### Effects of *C. intybus and N. sativa* aqueous extract on hematological parameters in alloxan-induced diabetic rabbits

RBC, WBC, Neutrophil, and PCV were decreased while other hematological parameters were increased in alloxan-induced diabetic rabbits. After four weeks of study, the result shows that the oral treatment of *C. intybus, N. sativa*, or their mixture significantly (*p* < 0.05) increased the RBC, WBC, neutrophil, and PCV% as compared with the positive control group. While the other hematological parameters were significantly (*p* < 0.05) decreased in treatment groups as compared with a positive control group. However, the effect of plant material on these parameters in diabetic animals was similar to standard drug-treated diabetic animals (Table 5).

**Table 5.** Effects of *Cichorium intybus* and *Nigella sativa* Aqueous Extract on Hematological Parameters in Alloxan-Induced Diabetic Rabbits.

| Parameters | I. Negative control | II. Positive control (diabetes-induced) | III. Diabetic + Metformin | IV. Diabetic + *Cichorium intybus* | V. Diabetic + *Nigella sativa* | VI. Diabetic + combination of plants |
| --- | --- | --- | --- | --- | --- | --- |
| RBC (x 10<sup>6</sup>/μl) | 2.26±1.08<sup>b</sup> | 5.89±1.04<sup>a</sup> | 5.00±1.29<sup>ca</sup> | 5.72±1.40<sup>ca</sup> | 4.96±1.1<sup>ca</sup> | 5.65±1.16<sup>ca</sup> |
| WBC (x 10<sup>3</sup>/μl) | 2.28±0.78<sup>b</sup> | 8.54±1.60<sup>a</sup> | 4.86±1.49<sup>c</sup> | 6.16±1.41<sup>c</sup> | 5.36±1.3<sup>c</sup> | 5.76±0.98<sup>c</sup> |
| Basophils % | 11.74±1.31<sup>b</sup> | 4.90±1.06<sup>a</sup> | 5.88±1.23<sup>ca</sup> | 7.10±2.45<sup>ca</sup> | 6.68±1.2<sup>ca</sup> | 5.50±1.31<sup>ca</sup> |
| Neutrophil % | 17.4±8.38<sup>b</sup> | 45.46±5.23<sup>a</sup> | 33.60±3.79<sup>c</sup> | 36.86±5.10<sup>c</sup> | 38.92±3.7<sup>c</sup> | 35.76±3.20<sup>c</sup> |
| Eosinophil % | 6.5±1.48<sup>b</sup> | 2.22±0.74<sup>a</sup> | 3.25±0.87<sup>ca</sup> | 3.74±0.97<sup>ca</sup> | 3.86±.89<sup>ca</sup> | 4.14±1.09<sup>ca</sup> |
| Monocytes % | 22.46±1.98<sup>b</sup> | 8.26±1.53<sup>a</sup> | 16.06±1.36<sup>c</sup> | 17.64±.097<sup>c</sup> | 15.26±.99<sup>c</sup> | 16.96±1.20<sup>c</sup> |
| Lymphocytes % | 69.72±3.03<sup>b</sup> | 42.02±3.30<sup>a</sup> | 51.92±5.46<sup>ca</sup> | 53.74±3.5<sup>ca</sup> | 55.28±4.3<sup>ca</sup> | 58.10±2.71<sup>ca</sup> |
| PCV % | 26.2±2.15<sup>b</sup> | 38.00±4.42<sup>a</sup> | 34.60±1.85<sup>ca</sup> | 30.60±7.08<sup>ca</sup> | 31.6±5.81<sup>ca</sup> | 29.80±5.26<sup>ca</sup> |

Each value represents the mean of 5 rabbits ± SD.
a: Significant difference between normal control and diabetic groups
b: Significant difference between diabetic untreated and diabetic treated groups
c: Non-significant difference between diabetic animals treated with standard drugs and diabetic animals treated with plant extract
ca: Non-significant difference among normal control animals and diabetic animals treated with standard drug or diabetic animals treated with plant extract

### Effects of *C. intybus and N. sativa* aqueous extract on urea and creatinine levels in alloxan induced diabetic rabbits

Creatinine and urea levels were increased in diabetic animals. At the end of the experiment, the present study result shows that urea and creatinine mean values were significantly (*p* < 0.05) decreased in *N. sativa, C. intybus*, or their mixture-treated diabetic rabbits when compared with positive control rabbits. However, the standard drug effect on urea and creatinine levels was slightly more efficient as compared with plant material (Table 6).

**Table 6.** Effects of *Cichorium intybus* and *Nigella sativa* Aqueous Extract on Urea and Creatinine Levels in Alloxan-Induced Diabetic Rabbits.

| Parameters | I. Negative control | II. Positive control (diabetes-induced) | III. Diabetic + Metformin | IV. Diabetic + *Cichorium intybus* | V. Diabetic + *Nigella sativa* | VI. Diabetic + combination of plants |
| --- | --- | --- | --- | --- | --- | --- |
| Urea (mg/dl) | 18.0± 4.24<sup>a</sup> | 124.2±7.22<sup>d</sup> | 43.8±9.1<sup>b</sup> | 60.4±6.15<sup>c</sup> | 68±8.55<sup>c</sup> | 64.4±11.53<sup>c</sup> |
| Creatinine (mg/dl) | 0.97 ±0.16<sup>a</sup> | 1.96 ±0.26<sup>d</sup> | 1.76±0.24<sup>b</sup> | 1.01 ±0.58<sup>c</sup> | 1.04±0.19<sup>c</sup> | 1.06±0.31<sup>c</sup> |

Each value represents the mean of 5 rabbits ± SD.
a: Significant difference between normal control and diabetic groups
b: Significant difference between diabetic untreated and diabetic treated groups
c: Non-significant difference between diabetic animals treated with standard drugs and diabetic animals treated with plant extract
d: Mean value of creatinine in negative control animals

### Effects of *C. intybus and N. sativa* aqueous extract on *AST, ALT, and ALP* enzyme levels in alloxan-induced diabetic rabbits

AST, ALT, and ALP values were significantly raised in diabetic animals as compared with normal levels. The result shows a significant (*p* < 0.05) decrease in the mean values of these enzymes in diabetic rabbits treated with *N. sativa, C. intybus*, or their mixture as compared to the positive control group. While slight difference in the mean values of these enzymes was shown in different treatment groups as compared with standard drugs. Moreover, the effects of plant material on these enzymes in different treatment groups were almost the same (Table 7).

**Table 7.** Effects of *Cichorium intybus* and *Nigella sativa* Aqueous Extract on AST, ALT, and ALP Enzyme Levels in Alloxan-Induced Diabetic Rabbits.

| Parameters | I. Negative control | II. Positive control (diabetes-induced) | III. Diabetic + Metformin | IV. Diabetic + *Cichorium intybus* | V. Diabetic + *Nigella sativa* | VI. Diabetic + combination of plants |
| --- | --- | --- | --- | --- | --- | --- |
| AST (U/L) | 62.6±15.5<sup>a</sup> | 127.4±10.3<sup>c</sup> | 90 ±19.7<sup>b</sup> | 94.8±8.1<sup>b</sup> | 101.2±8.5<sup>b</sup> | 99.6±8.3<sup>b</sup> |
| ALT (U/L) | 29.1±5.2<sup>a</sup> | 96.4±7.2<sup>c</sup> | 92.2±4.4<sup>b</sup> | 78.4±5.1<sup>b</sup> | 84.6±5.6<sup>b</sup> | 85.4±7.1<sup>b</sup> |
| ALP (U/L) | 95.8±32.8<sup>a</sup> | 149.4±9.4<sup>c</sup> | 126.4±10.9<sup>b</sup> | 117.2±11.9<sup>ab</sup> | 113.8±11.1<sup>ab</sup> | 114.6±12.4<sup>ab</sup> |

Each value represents the mean of 5 rabbits ± SD.
a: Significant difference between normal control and diabetic groups
b: non-significant difference between diabetic animals treated with standard drugs and diabetic animals treated with plant extract
c: Significant difference between diabetic untreated and diabetic treated groups
ab: Non-significant difference among normal control animals and diabetic animals treated with plant extract

## DISCUSSION

In our study we observed that *N. sativa* and *C. intybus* plant extract provides significant hypoglycemic effects in diabetic animals. Our results show that the mean blood glucose values were decreased gradually in diabetic animals treated with *N. sativa* and *C. intybus*. Diabetic animals treated with *N. sativa and C. intybus* or their mixture showed a significant (*p* < 0.05) decreased in blood sugar levels as compared with positive control animals. These results were similar to studies that observed a hypoglycemic effect of these plants on different treatment days in alloxan-induced diabetic rabbits (Hassan & Yousef, 2010; Gorjipour et al., 2017). Research also shows that herbal medicine and their active ingredients have potential hypoglycemic effect in experimental animals (Kanter 2009; Rub et al., 2014). Body weight and blood glucose levels of animals are indicators of normalized metabolism, particularly carbohydrate metabolism which is affected primarily in diabetes.

The results show that the mean body weight gain values of treatment groups were significantly (*p* < 0.05) increased as compared with the positive control group. While, a slight difference in these values was shown in *N. sativa, C. intybus*, or their mixture-treated diabetic animals as compared with standard drugs. Our research follows (Goth 1991; Rezagholizadeh et al., 2016), which noticed a non-significant difference in body weight gain in animals treated with plant material as compared with standard treated animals.

The positive effects of *N. sativa* and *C. intybus* in treatment animals could be due to the efficacy of these plants in restraining angiogenesis in adipose tissue and reducing differentiation of preadipocytes (Djilani et al., 2011).

Previous observations showed that the decrease in serum insulin levels in alloxan-induced diabetic rabbits was due to the cytotoxic effect of alloxan monohydrate on b-cells of pancreases (Judžentienė & Būdienė, 2008). The current study result shows that *N. Sativa, C. intybus*, or their mixture treatment significantly (*p* < 0.05) increased the serum insulin level in diabetic groups as compared with the positive control group. Our study results were similar to (Zareen et al., 2008), who observed that herbal medicine increased the serum insulin level in diabetic animals. Meanwhile, the effect of plant material to increase serum insulin levels in diabetic groups was comparable to metformin-treated diabetic group.

RBC, WBC, Neutrophil, and Pack cell volume % were decreased while other hematological parameters were increased in alloxan-induced diabetic animals. The increased heart rate was apparently due to the enhanced sympathetic system of diabetic animals, produced by diabetes-induced anemia (Al-Okbi et al., 2013). (Verma et al., 2013), investigated the cardiovascular effects of plant extracts on diabetic animals. Their results were similar to our findings.

The present study result shows that diabetic animals treated with plant material and standard drug resulted in a significant (*p* < 0.05) increased in the RBC, WBC, Neutrophil, and PCV% as compared with positive control animals. While the other hematological parameters were significantly (*p* < 0.05) decreased in treatment animals as compared with positive control animals. These results were similar to (Kumar & Suresh, 2012), who observed that *N. sativa* and *C. intybus* treatment decreased heart rate and increased pack cell volume % to normal level in diabetic animals. Hence, the reduced heart rate was may be due to a normalized PCV% and other hematological parameters in these animals.

The current study shows that urea and creatinine levels were increased significantly in alloxan-induced diabetic rabbits. A similar study was conducted by (Ozkol et al., 2013), which observed the increased mean values of urea and creatinine concentration, and indicated that renal function was impaired in diabetes. The breakdown of amino acids during gluconeogenesis results in increased urea production and negative nitrogen balance in diabetic animals (Tuorkey 2017). At the end of the experiment, the result shows that urea and creatinine mean values were significantly (*p* < 0.05) decreased in *N. Sativa, C. intybus*, or their mixture-treated diabetic rabbits when compared with positive control rabbits. However, the standard drug effect on urea and creatinine levels was slightly more efficient as compared with plant material.

AST, ALT and ALP values in diabetic rabbits were higher as compared to normal levels. The increased values of these enzymes in the blood are due to the leakage of these enzymes from the liver cytosol into the bloodstream (Abdelrazek et al., 2018), and show that alloxan monohydrate has a hepatotoxic impact. The present study result shows a significant (*p* < 0.05) decrease in the mean values of these enzymes in diabetic rabbits treated with plant material or their mixture as compared to the positive control group. While slight difference in the mean values of these enzymes was shown in different treatment groups as compared with standard drugs. Moreover, the effects of plant material on these enzyme values in different treatment groups were almost the same. These findings were similar to (Zaman 2011; Najmi et al., 2008), who reported that treatment of diabetic animals with these plants for a longer time significantly decreased enzyme levels to near normal values.

## CONCLUSION

It is reasonable to conclude that *N. sativa* seed and *C. intybus* leaf extracts possess strong anti-diabetic properties. Indeed, the results of the effects of studied plants are comparable with the anti-diabetic effect of standard drug metformin. The studied plant extract treatment maintained the body weight, decreased the urea and creatinine levels, and increased the activity of liver enzymes in diabetic rabbits. It could be well anticipated that both of these plants have a protectant effect against kidney and liver damage in diabetics. Further studies are needed to trace out and exploit the active ingredients of these two promising plant extracts.

## Ethical statement

There is no statement against the values of animals used in the experiment. The animals were maintained following “Principals of Laboratory Animal Care’’. With the approval of the Ethic Committee of the University of Veterinary & Animal Sciences (UVAS), Lahore (No. CVAS/13545 dated 07-01-2021) and the Directorate of Advanced Studies (DAS) of the same university (DAS/536 dated 15-06-2021).

## Acknowledgment

This study was supported by the National Research Program for Universities, Grant/Award Number: 20-14678/NRPU/R&D/HEC/2021.

